# Antiviral humoral immunity against SARS-CoV-2 Omicron subvariants induced by XBB.1.5 monovalent vaccine in infection-naïve and XBB-infected individuals

**DOI:** 10.1101/2023.11.29.569330

**Authors:** Yusuke Kosugi, Yu Kaku, Alfredo A Hinay, Ziyi Guo, Keiya Uriu, Minoru Kihara, Fumitake Saito, Yoshifumi Uwamino, Jin Kuramochi, Kotaro Shirakawa, Akifumi Takaori-Kondo, The Genotype to Phenotype Japan (G2P-Japan) Consortium, Kei Sato

## Abstract

To control infection with SARS-CoV-2 Omicron XBB subvariants, the XBB.1.5 monovalent mRNA vaccine has been available since September 2023. However, we have found that natural infection with XBB subvariants, including XBB.1.5, does not efficiently induce humoral immunity against the infecting XBB subvariants. These observations raise the possibility that the XBB.1.5 monovalent vaccine may not be able to efficiently induce humoral immunity against emerging SARS-CoV-2 variants, including a variety of XBB subvariants (XBB.1.5, XBB.1.16, XBB.2.3, EG.5.1 and HK.3) as well as BA.2.86. To address this possibility, we collected two types of sera from individuals vaccinated with the XBB.1.5 vaccine; those who had not been previously infected with SARS-CoV-2 and those who had been infected with XBB subvariants prior to XBB.1.5 vaccination. We collected sera before and 3-4 weeks after vaccination, and then performed a neutralization assay using these sera and pseudoviruses.

## Text

To control infection with SARS-CoV-2 Omicron XBB subvariants, the XBB.1.5 monovalent mRNA vaccine has been available since September 2023. However, we have found that natural infection with XBB subvariants, including XBB.1.5, does not efficiently induce humoral immunity against the infecting XBB subvariants.^1-3^ These observations raise the possibility that the XBB.1.5 monovalent vaccine may not be able to efficiently induce humoral immunity against emerging SARS-CoV-2 variants, including a variety of XBB subvariants (XBB.1.5, XBB.1.16, XBB.2.3, EG.5.1 and HK.3) as well as BA.2.86. To address this possibility, we collected two types of sera from individuals vaccinated with the XBB.1.5 vaccine; those who had not been previously infected with SARS-CoV-2 (N=9; **Figure A**) and those who had been infected with XBB subvariants prior to XBB.1.5 vaccination (N=10; **Figure B**). We collected sera before and 3-4 weeks after vaccination, and then performed a neutralization assay using these sera and pseudoviruses. As expected, XBB.1.5 vaccine sera with prior XBB infection efficiently (1.8-to 3.6-fold) boosted antiviral humoral immunity against all variants tested with statistical significance (**Figure B**). Importantly, in the case of the XBB.1.5 vaccine sera without prior infection, XBB.1.5 vaccine also induced efficiently antiviral activity (2.1-to 3.9-fold) against all variants tested with statistical significance (**Figure A**). These observations suggest that a single dose of XBB.1.5 monovalent vaccine potentially induces antiviral humoral immunity against XBB subvariants as well as BA.2.86 without prior infection.

**Figure.**
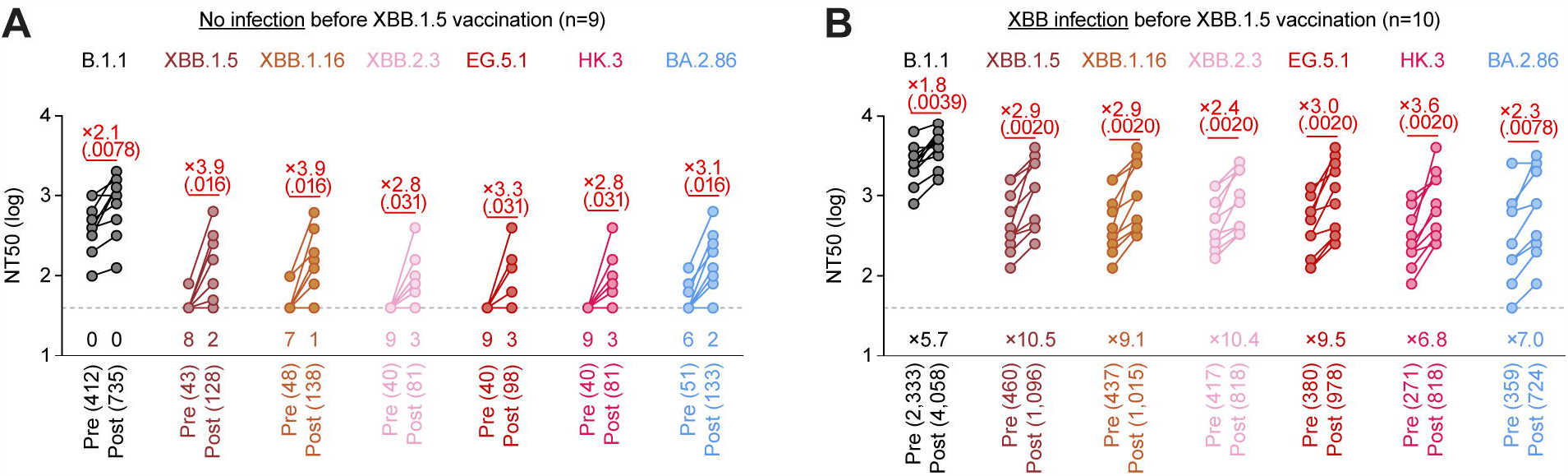
The neutralization activity induced by XBB.1.5 monovalent vaccine. Assays were performed with pseudoviruses harboring the S proteins of B.1.1, XBB.1.5, XBB.1.16, XBB.2.3, EG.5.1, HK.3 and BA.2.86. The following two sera were used: (**A**) vaccinated sera from fully vaccinated individuals who had not been infected (‘<u>no infection</u> before XBB.1.5 vaccination’, 9 donors), (**B**) vaccinated sera from fully vaccinated individuals who had been infected with XBB subvariants (after June 2023) (‘<u>XBB infection</u> before XBB.1.5 vaccination’, 10 donors). Sera were collected before vaccination (‘Pre’) and 3-4 weeks after XBB.1.5 monovalent vaccination (‘Post’). Assays for each serum sample were performed in triplicate to determine the 50% neutralization titer (NT_50_). Each dot represents the NT_50_ value for each donor, and the NT_50_ values for the same donor between pre- and post-vaccination are connected by a line. The number in parenthesis below the graph indicates the geometric mean of the NT_50_ value. The horizontal dashed line indicates the detection limit (40-fold dilution). In **A**, the number of the serum donors with the NT_50_ values below the detection limit is shown in the figure (below the horizontal dashed line). In **B**, the number with “X” (below the horizontal dashed line) indicates the fold change of the NT_50_ value of the pre-vaccination of the XBB-infected cohort samples (**B**) compared to that of the infection-naïve cohort samples (**A**). Statistically significant differences between pre-and post-vaccination were determined by two-sided Wilcoxon signed-rank tests and are shown in red parentheses. The fold change of the reciprocal NT_50_ is calculated between pre- and post-vaccination and is shown in red. Background information on the vaccinated donors is summarized in **Table S1**.

The induction efficiency of neutralizing activity was comparable between the infection-naïve cohort (**Figure A**) and the XBB-infected cohort (**Figure B**). However, in sera collected prior to XBB.1.5 vaccination, the 50% neutralization titer of sera from the XBB-infected cohort was 5.7-to 10.4-fold higher than that of sera from the infection-naïve cohort (**Figure**). In addition, although all pre-vaccination sera of XBB-infected cohort exhibited antiviral activity against all variants tested, some individuals vaccinated with XBB.1.5 vaccine without natural infection showed no antiviral activity against XBB.1.5 (N=2), XBB.1.16 (N=1), XBB.2.3 (N=3), EG.5.1 (N=3), HK.3 (N=3) and BA.2.86 (N=2). Taken together, these results suggest that a single dose of XBB.1.5 monovalent vaccine may not be sufficient to induce effective antiviral humoral immunity in infection-naïve individuals and that a booster dose of XBB.1.5 monovalent vaccine may be required in some cases.

## Grants

Supported in part by AMED SCARDA Japan Initiative for World-leading Vaccine Research and Development Centers “UTOPIA” (JP223fa627001, to Kei Sato), AMED SCARDA Program on R&D of new generation vaccine including new modality application (JP223fa727002, to Kei Sato); AMED Research Program on Emerging and Re-emerging Infectious Diseases (JP22fk0108146, to Kei Sato; JP21fk0108494 to G2P-Japan Consortium, Kei Sato; JP21fk0108425, to Kei Sato; JP21fk0108432, to Kei Sato; JP22fk0108511, to G2P-Japan Consortium and Kei Sato; JP22fk0108516, to Kei Sato; JP22fk0108506, to Kei Sato; JP22fk0108534, to Kei Sato); AMED Research Program on HIV/AIDS (JP22fk0410039, to Kei Sato); JST CREST (JPMJCR20H4, to Kei Sato); JSPS Core-to-Core Program (A. Advanced Research Networks) (JPJSCCA20190008, Kei Sato); JSPS Research Fellow DC2 (22J11578, to Keiya Uriu); JSPS Research Fellow DC1 (23KJ0710, to Yusuke Kosugi); The Tokyo Biochemical Research Foundation (to Kei Sato); and The Mitsubishi Foundation (to Kei Sato).

## References

1. Uriu K, Ito J, Kosugi Y, et al. Transmissibility, infectivity, and immune evasion of the SARS-CoV-2 BA.2.86 variant. Lancet Infect Dis 2023; 23(11): e460–e1.

2. Kosugi Y, Plianchaisuk A, Putri O, et al. Virological characteristics of the SARS-CoV-2 Omicron HK.3 variant harboring the “FLip” substitution. BioRxiv 2023: doi: 10.1101/2023.11.14.566985.

3. Kaku Y, Kosugi Y, Uriu K, et al. Antiviral efficacy of the SARS-CoV-2 XBB breakthrough infection sera against omicron subvariants including EG.5. Lancet Infect Dis 2023; 23(10): e395–e6.

