## Supplementary Appendix for "Antiviral humoral immunity against SARS-CoV-2 Omicron subvariants induced by XBB.1.5 monovalent vaccine in infection-naïve and XBB-infected individuals"

**Table of Contents**

**Contents** Page


**Materials and Methods** 2-3
Ethics statement
Human serum collection
Plasmid construction
Cell culture
Pseudovirus preparation
Neutralization assay
**Table S1.** Human sera used in this study 4
**Table S2.** Primers used in this study 5
**Consortia** 6
**Acknowledgments** 7
**Supplemental References** 8

### Materials and Methods

#### Ethics statement

All protocols involving specimens from human subjects recruited at The Institute of Medical Science, Kyoto University, Interpark Kuramochi Clinic, Namikibashi Clinic and Wakaba Clinic was reviewed and approved by the Institutional Review Boards of The Institute of Medical Science (approval IDs: 2021-1-0416 and 2022-29-0915), Kyoto University (approval ID: G1309), Interpark Kuramochi Clinic (approval ID: G2021-004), Namikibashi Clinic and Wakaba Clinic (approval ID: 2022-29-0915), respectively. All human subjects provided written informed consent. All protocols for the use of human specimens were reviewed and approved by the Institutional Review Boards of The Institute of Medical Science, The University of Tokyo (approval IDs: 2021-1-0416, 2021-18-0617 and 2022-29-0915).

#### Human serum collection

XBB.1.5 monovalent vaccine sera from fully vaccinated individuals who had not been infected (nine donors. Average age, 66.6; range, 51–89; 33.3% male) and those from fully vaccinated individuals who had been infected with XBB subvariants (ten donors. Average age, 48.3; range, 32–61; 70% male) were collected before vaccination and three weeks (20–28 days) after vaccination. SARS-CoV-2 variants were genotyped in four out of the ten infected individuals.[1-3] In six out of the ten infected individuals, the infecting variant was classified as XBB subvariants without genotyping, because after June 2023, only XBB subvariants were detected in Japan (**Table S1**). The SARS-CoV-2 variants were identified as previously described.[4-6] Sera were inactivated at 56°C for 30 minutes and stored at –80°C until use. The details of the convalescent sera are summarized in **Table S1**.

#### Plasmid construction

Plasmids expressing the SARS-CoV-2 spike (S) proteins of B.1.1 (the parental D614G-bearing variant), Omicron XBB.1.5, XBB.1.16, EG.5.1, HK.3 and BA.2.86 were prepared in our previous studies.[1-3,7-9] Plasmids expressing the S protein of XBB.2.3 was generated by site-directed overlap extension PCR using pC-SARS2-S XBB.1.5[7] as the template and the primers listed in **Table S2**. The resulting PCR fragment was subcloned into the KpnI-NotI site of the pCAGGS vector[10] using In-Fusion HD Cloning Kit (Takara, Cat# Z9650N). Nucleotide sequences were determined by DNA sequencing services (Eurofins), and the sequence data were analyzed by SnapGene software v6.1.1 (www.snapgene.com).

#### Cell culture

Lenti-X 293T cells (Takara, Cat# 632180) and HOS-ACE2/TMPRSS2 cells (kindly provided by Dr. Kenzo Tokunaga), a derivative of HOS cells (a human osteosarcoma cell line; ATCC CRL-1543) stably expressing human ACE2 and TMPRSS2,[9,11] were maintained in Dulbecco's modified

Eagle's medium (DMEM) (high glucose) (Wako, Cat# 044- 29765) containing 10% fetal bovine serum (Sigma-Aldrich Cat# 172012-500ML), 100 units penicillin and 100 ug/ml streptomycin (Sigma-Aldrich, Cat# P4333-100ML).

**Pseudovirus preparation**

Pseudoviruses were prepared as previously described.[8] Briefly, lentivirus (HIV-1)-based, luciferase-expressing reporter viruses were pseudotyped with the SARS-CoV-2 S proteins. One prior day of transfection, the LentiX-293T cells were seeded at a density of 2 × $10^6$ cells. The LentiX-293T cells were cotransfected with 1 µg psPAX2-IN/HiBiT (a packaging plasmid encoding the HiBiT-tag-fused integrase[12]), 1 µg pWPI-Luc2 (a reporter plasmid encoding a firefly luciferase gene[12]) and 500 ng plasmids expressing parental S or its derivatives using TransIT-293 transfection reagent (Mirus, Cat# MIR2704) according to the manufacturer's protocol. Two days post transfection, the culture supernatants were harvested and filtrated. The pseudoviruses were stored at –80°C until use.

**Neutralization assay**

Neutralization assays were performed as previously described.[8] The SARS-CoV-2 spike pseudoviruses (counting ~100,000 relative light units) were incubated with serially diluted (40-fold to 29,160-fold dilution at the final concentration) heat-inactivated sera at 37°C for 1 hour. Pseudoviruses without sera were included as controls. Then, 20 µl mixture of pseudovirus and serum was added to HOS-ACE2/TMPRSS2 cells (10,000 cells/100 µl) in a 96-well white plate. Two days post infection, the infected cells were lysed with a Bright-Glo luciferase assay system (Promega, Cat# E2620), and the luminescent signal was measured using a GloMax explorer multimode microplate reader 3500 (Promega). The assay of each serum sample was performed in triplicate, and the 50% neutralization titer was calculated using Prism 9 (GraphPad Software).

**Table S1. Human sera used in this study**

| Donor ID | Sex | Age | Date of 1st vaccination (YYYY-MM-DD) | Date of 2nd vaccination (YYYY-MM-DD) | Date of 3rd vaccination (YYYY-MM-DD) | Date of 4th vaccination (YYYY-MM-DD) | Date of 5th vaccination (YYYY-MM-DD) | Date of 6th vaccination (YYYY-MM-DD) | Date of sampling (before vaccination) (YYYY-MM-DD) | Date of XBB.1.5 vaccination (YYYY-MM-DD) | Date of sampling (after vaccination) (YYYY-MM-DD) | Time interval between vaccination and the second sampling | Prior infection? | variant |
|---|---|---|---|---|---|---|---|---|---|---|---|---|---|---|
| 5165 | Female | 89 | 2021-05-29 (P) | 2021-06-21 (P) | 2022-02-16 (P) | 2022-07-17 (P) | 2022-11-27 (PBA4/5) | 2023-05-20 (PBA4/5) | 2023-09-29 | 2023-09-29 (XBB1.5) | 2023-10-26 | 27 | No | |
| 5166 | Male | 77 | 2021-06-05 (P) | 2021-07-04 (P) | 2022-03-05 (P) | 2022-08-06 (P) | 2022-11-20 (PBA4/5) | 2023-05-20 (PBA4/5) | 2023-09-29 | 2023-09-29 (XBB1.5) | 2023-10-21 | 22 | No | |
| 6783 | Male | 57 | 2021-06-23 (M) | 2021-07-21 (M) | 2022-02-11 (M) | 2022-10-15 (MBA1) | | | 2023-10-03 | 2023-10-03 (XBB1.5) | 2023-10-23 | 20 | No | |
| 2292 | Female | 84 | 2021-06-01 (P) | 2021-06-22 (P) | 2022-02-25 (P) | 2022-07-28 (P) | 2022-11-07 (PBA4/5) | 2023-05-30 (PBA4/5) | 2023-10-05 | 2023-10-05 (XBB1.5) | 2023-10-26 | 21 | No | |
| 6858 | Female | 62 | 2021-07-18 (P) | 2021-08-11 (P) | 2022-02-27 (M) | 2022-07-30 (P) | 2022-11-20 (PBA4/5) | | 2023-10-07 | 2023-10-07 (XBB1.5) | 2023-10-28 | 21 | No | |
| 192 | Female | 64 | 2021-04-22 (P) | 2021-05-13 (P) | 2022-01-15 (P) | 2022-07-16 (P) | 2022-11-24 (PBA4/5) | 2023-05-25 (PBA4/5) | 2023-10-02 | 2023-10-26 (PXBB1.5) | 2023-11-20 | 25 | No | |
| 1700 | Female | 53 | 2021-04-21 (P) | 2021-05-12 (P) | 2022-01-15 (P) | 2022-07-13 (P) | 2022-11-30 (PBA4/5) | 2023-06-23 (MBA4/5) | 2023-09-29 | 2023-10-18 (PXBB1.5) | 2023-11-13 | 26 | No | |
| 5555 | Female | 51 | 2021-07-14 (P) | 2021-08-14 (P) | 2022-02-22 (P) | 2022-07-23 (P) | 2022-12-03 (PBA4/5) | 2023-05-11 (PBA4/5) | 2023-09-30 | 2023-10-21 (PXBB1.5) | 2023-11-15 | 25 | No | |
| 5986 | Male | 62 | 2021-07-24 (P) | 2021-08-14 (P) | 2022-03-12 (M) | 2022-08-27 (M) | 2022-12-24 (PBA4/5) | 2023-06-03 (PBA4/5) | 2023-09-25 | 2023-10-21 (PXBB1.5) | 2023-11-13 | 23 | No | |
| KS | Male | 41 | 2021-06-17 (P) | 2021-07-07 (P) | 2022-03-28 (M) | 2022-10-27 (MBA.5) | | | 2023-09-19 | 2023-09-20 (XBB1.5) | 2023-10-14 | 24 | Yes (2023-06-29) | XBB.1.9 |
| KY | Female | 53 | 2021-08-18 (P) | 2021-09-08 (P) | 2022-04-13 (P) | 2022-10-21 (P) | | | 2023-09-25 | 2023-09-27 (XBB1.5) | 2023-10-20 | 23 | Yes (2023-07-24) | XBB.1.16 |
| KK | Male | 56 | 2021-07-16 (P) | 2021-08-06 (P) | 2022-03-03 (M) | 2022-08-09 (P) | | | 2023-09-25 | 2023-09-29 (XBB1.5) | 2023-10-24 | 25 | Yes (2023-07-17) | EG.5 |
| 2345 | Female | 52 | 2021-07-11 (P) | 2021-08-01 (P) | 2022-03-10 (P) | 2022-10-22 (PBA1) | | | 2023-10-06 | 2023-10-06 (XBB1.5) | 2023-10-27 | 21 | Yes (2023-07) | NA |
| 80 | Male | 61 | 2021-05-10 (P) | 2021-05-31 (P) | 2022-01-24 (M) | 2022-07-22 (BA4/5) | 2022-11-12 (BA4/5) | 2023-06-03 (BA4/5) | 2023-09-29 | 2023-09-30 (XBB1.5) | 2023-10-24 | 24 | Yes (2023-07) | NA |
| 90 | Male | 47 | 2021-05-11 (P) | 2021-06-02 (P) | 2022-01-25 (M) | 2022-08-02 (BA4/5) | 2022-11-09 (BA4/5) | | 2023-10-10 | 2023-10-11 (XBB1.5) | 2023-11-01 | 21 | Yes (2023-07) | NA |
| 100 | Male | 32 | 2021-05-18 (P) | 2021-06-08 (P) | 2022-01-31 (M) | 2022-07-17 (MBA4/5) | 2022-11-15 (PBA4/5) | 2023-05-27 (MBA4/5) | 2023-10-11 | 2023-10-14 (XBB1.5) | 2023-11-06 | 23 | Yes (2023-06) | NA |
| 286691 | Male | 49 | 2021-06-01(P) | 2021-06-22 (P) | 2022-03-21 (M) | | | | 2023-10-19 | 2023-10-19 (PXBB1.5) | 2023-11-16 | 28 | Yes (2023-08-22) | XBB.1.9 |
| 2439306 | Male | 43 | 2021-03-08 (P) | 2021-03-29 (P) | 2021-12-20 (P) | 2022-08-26 (M) | 2022-11-30 (M) | | 2023-10-23 | 2023-10-23 (PXBB1.5) | 2023-11-20 | 28 | Yes (2023-09-08) | NA |
| 4177 | Female | 49 | 2021-04-24 (P) | 2021-05-15 (P) | 2022-01-22 (P) | 2022-07-09 (P) | 2022-12-03 (PBA4/5) | 2023-06-13 (MBA4/5) | 2023-09-29 | 2023-10-14 (PXBB1.5) | 2023-11-10 | 27 | Yes (2023-08-29) | NA |

NA, not applicable.

P, Pfizer-BioNTech; M, Moderna

**Table S2. Primers used in this study**

| Primer name | Primer sequence (5'-to-3') | Purpose |
|---|---|---|
| Omicron universal Fw | cactatagggcgaattgggtaccatgtttgtgttcctggt | Preparation of S expression plasmid |
| BA.2 WT Rv | agctccaccgcggtggcggccgctcaggtgtagtgcagtttca | Preparation of S expression plasmid |
| V252G_D253G_Fw | tcctacctgacacctGGAGGAtcctcctctggctgg | Preparation of S expression plasmid |
| V252G_D253G_Rv | ccagccagaggaggaTCCTCCaggtgtcaggtagga | Preparation of S expression plasmid |
| P521S_Fw | gaactgctccatgccTCTgccacagtgtgtgg | Preparation of S expression plasmid |
| P521S_Rv | ccacacactgtggcAGAggcatggagcagttc | Preparation of S expression plasmid |

**Consortia**

**The Genotype to Phenotype Japan (G2P-Japan) Consortium**

**The Institute of Medical Science, The University of Tokyo, Japan**

Jumpei Ito, Naoko Misawa, Arnon Plianchaisuk, Kaoru Usui, Wilaiporn Saikruang, Shigeru Fujita, Jarel Elgin Mendoza Tolentino, Luo Chen, Lin Pan, Mai Suganami, Mika Chiba, Ryo Yoshimura, Kyoko Yasuda, Keiko Iida, Adam Patrick Strange, Naomi Ohsumi, Shiho Tanaka, Kaho Okumura

**Hokkaido University, Japan**

Takasuke Fukuhara, Tomokazu Tamura, Rigel Suzuki, Saori Suzuki, Hayato Ito, Keita Matsuno, Hirofumi Sawa, Naganori Nao, Shinya Tanaka, Masumi Tsuda, Lei Wang, Yoshikata Oda, Zannatul Ferdous, Kenji Shishido

**Tokyo Metropolitan Institute of Public Health**

Kazuhisa Yoshimura, Kenji Sadamasu, Mami Nagashima, Hiroyuki Asakura

**Tokai University, Japan**

So Nakagawa

**Kyoto University, Japan**

Kayoko Nagata, Ryosuke Nomura, Yoshihito Horisawa, Yusuke Tashiro, Yugo Kawai, Kazuo Takayama, Rina Hashimoto, Sayaka Deguchi, Yukio Watanabe, Ayaka Sakamoto, Naoko Yasuhara, Takao Hashiguchi, Tateki Suzuki, Kanako Kimura, Jiei Sasaki, Yukari Nakajima, Hisano Yajima

**Hiroshima University, Japan**

Takashi Irie, Ryoko Kawabata

**Kyushu University, Japan**

Kaori Tabata

**Kumamoto University, Japan**

Terumasa Ikeda, Hesham Nasser, Ryo Shimizu, MST Monira Begum, Michael Jonathan, Yuka Mugita, Otowa Takahashi, Kimiko Ichihara, Takamasa Ueno, Chihiro Motozono, Mako Toyoda

**University of Miyazaki, Japan**

Akatsuki Saito, Maya Shofa, Yuki Shibatani, Tomoko Nishiuchi

**Charles University, Czechia**

Jiri Zahradnik, Prokopios Andrikopoulos, Miguel Padilla-Blanco, Aditi Konar

### Acknowledgments

We would like to thank all members of The Genotype to Phenotype Japan (G2P-Japan) Consortium. We thank Dr. Kenzo Tokunaga (National Institute of Infectious Diseases, Japan) for sharing materials.

### Supplementary References


1. Uriu K, Ito J, Kosugi Y, et al. Transmissibility, infectivity, and immune evasion of the SARS-CoV-2 BA.2.86 variant. *Lancet Infect Dis* 2023; **23**(11): e460-e1.
2. Kosugi Y, Plianchaisuk A, Putri O, et al. Virological characteristics of the SARS-CoV-2 Omicron HK.3 variant harboring the "FLip" substitution. *BioRxiv* 2023: doi: https://doi.org/10.1101/2023.11.14.566985.
3. Kaku Y, Kosugi Y, Uriu K, et al. Antiviral efficacy of the SARS-CoV-2 XBB breakthrough infection sera against omicron subvariants including EG.5. *Lancet Infect Dis* 2023; **23**(10): e395-e6.
4. Yamasoba D, Kimura I, Nasser H, et al. Virological characteristics of the SARS-CoV-2 Omicron BA.2 spike. *Cell* 2022; **185**(12): 2103-15.e19.
5. Kimura I, Yamasoba D, Tamura T, et al. Virological characteristics of the novel SARS-CoV-2 Omicron variants including BA.4 and BA.5. *Cell* 2022; **185**(21): 3992-4007.e16.
6. Saito A, Tamura T, Zahradnik J, et al. Virological characteristics of the SARS-CoV-2 Omicron BA.2.75 variant. *Cell Host Microbe* 2022; **30**(11): 1540–55.e15.
7. Uriu K, Ito J, Zahradnik J, et al. Enhanced transmissibility, infectivity, and immune resistance of the SARS-CoV-2 omicron XBB.1.5 variant. *Lancet Infect Dis* 2023; **23**(3): 280-1.
8. Yamasoba D, Uriu K, Plianchaisuk A, et al. Virological characteristics of the SARS-CoV-2 omicron XBB.1.16 variant. *Lancet Infect Dis* 2023; **23**(6): 655-6.
9. Ozono S, Zhang Y, Ode H, et al. SARS-CoV-2 D614G spike mutation increases entry efficiency with enhanced ACE2-binding affinity. *Nat Commun* 2021; **12**(1): 848.
10. Niwa H, Yamamura K, Miyazaki J. Efficient selection for high-expression transfectants with a novel eukaryotic vector. *Gene* 1991; **108**(2): 193-9.
11. Ferreira I, Kemp SA, Datir R, et al. SARS-CoV-2 B.1.617 mutations L452R and E484Q are not synergistic for antibody evasion. *J Infect Dis* 2021; **224**(6): 989-94.
12. Ozono S, Zhang Y, Tobiume M, Kishigami S, Tokunaga K. Super-rapid quantitation of the production of HIV-1 harboring a luminescent peptide tag. *J Biol Chem* 2020; **295**(37): 13023-30.
